# Variable organization of symbiont-containing tissue across planthoppers hosting different heritable endosymbionts

**DOI:** 10.1101/2022.12.06.519352

**Authors:** Anna Michalik, Diego C. Franco, Junchen Deng, Teresa Szklarzewicz, Michał Kobiałka, Adam Stroiński, Piotr Łukasik

**Affiliations:** Department of Developmental Biology and Morphology of Invertebrates, Institute of Zoology and Biomedical Research, Faculty of Biology, Jagiellonian University, Krakow, Poland; Institute of Environmental Sciences, Faculty of Biology, Jagiellonian University, Krakow, Poland; Museum and Institute of Zoology, Polish Academy of Sciences, Warsaw, Poland

**Keywords:** planthoppers, nutritional endosymbiosis, bacteriome, symbionts

## Abstract

Sap-feeding hemipteran insects live in associations with diverse heritable endosymbiotic bacteria and fungi that provide essential nutrients deficient in their diets. These symbionts typically reside in bacteriomes, dedicated organs made up of specialized cells termed bacteriocytes. The bacteriome organization varies between insect clades that are ancestrally associated with different microbes. As these symbioses evolve and additional microorganisms complement or replace the ancient associates, the organization of the symbiont-containing tissue becomes even more variable.

Planthoppers (Hemiptera: Fulgoromorpha) are ancestrally associated with bacterial symbionts *Sulcia* and *Vidania*, but in many of the planthopper lineages, these symbionts are now accompanied or have been replaced by other heritable bacteria (e.g., *Sodalis, Arsenophonus, Purcelliella*) or fungi. We know the identity of many of these microbes, but the symbiont distribution within the host tissues and the bacteriome organization have not been systematically studied using modern microscopy techniques.

Here, we combine light, fluorescence, and transmission electron microscopy with phylogenomic data to compare symbiont tissue distributions and the bacteriome organization across planthoppers representing 15 families. We identify and describe seven primary types of symbiont localization and seven types of the organization of the bacteriome. We show that *Sulcia* and *Vidania*, when present, occupy distinct bacteriomes distributed within the body cavity. The more recently acquired gammaproteobacterial and fungal symbionts generally occupy separate groups of cells organized into distinct bacteriomes or mycetomes, distinct from those with *Sulcia* and *Vidania*. They can also be localized in the cytoplasm of fat body cells. Alphaproteobacterial symbionts colonize a wider range of host body habitats: *Asaia*-like symbionts often colonize the host gut lumen, whereas *Wolbachia* and *Rickettsia* are usually scattered across tissues and cell types, including bacteriocytes containing other symbionts, bacteriome sheath, fat body cells, gut epithelium, as well as hemolymph. However, there are exceptions, including Gammaproteobacteria that share bacteriome with *Vidania*, or Alphaproteobacteria that colonize *Sulcia* cells.

We discuss how planthopper symbiont localization correlates with their acquisition and replacement patterns and the symbionts’ likely functions. We also discuss the evolutionary consequences, constraints, and significance of these findings.

## 1. Introduction

Intimate symbiotic relationships with microorganisms have played significant roles in the biology of many insects, influencing their nutrition, reproduction, development, protection against antagonists, and toxin resistance (Baumann, 2005; Flórez et al., 2015). Symbiosis with microbes is also among the crucial drivers of insects’ evolutionary diversification and adaptation to diverse environmental and biotic challenges associated with diverse global ecosystems (Feldhaar, 2011). Insect symbioses may involve both prokaryotic and eukaryotic microorganisms and differ in complexity and stability (Buchner, 1965; Douglas, 2016; Frago et al., 2020; McCutcheon et al., 2019). On one end of the spectrum, many insects seem to associate with transient microbes acquired from the environment every generation (Hammer and Moran, 2019). On the other end, symbioses involving ancient heritable nutritional microorganisms date back hundreds of millions of years and are characterized by extreme host-symbiont integration resulting mainly from metabolic dependence (Bennett and Moran, 2013; Moran et al., 1993). At the same time, other heritable nutritional symbioses, including those with fungi or some gammaproteobacteria, seem to be relatively recent (Husnik and McCutcheon, 2016; Matsuura et al., 2018; Michalik et al., 2021).

Among the most complex endosymbioses are symbiotic systems of Auchenorrhyncha - a suborder of hemipterans feeding almost exclusively on phloem or xylem plant saps. To overcome such a restricted and unbalanced diet, those insects have established mutualistic associations with diverse endosymbionts that provide them with essential amino acids and vitamins deficient in the sap (Douglas, 2016, 2009). Many Auchenorrhyncha harbor two heritable, nutritional co-primary bacterial symbionts, which complement each other in the supplementation of the insect’s diet. The ancestral symbiont of all Auchenorrhyncha is *Sulcia muelleri* (Bacteroidetes; hereafter *Sulcia*), which colonized the common ancestor of these insects ca. 300 mya and has co-diversified with the hosts since then (Moran et al., 2005). *Sulcia* is always accompanied by additional microbes. Depending on the Auchenorrhyncha superfamily, its ancestral co-symbionts include betaproteobacteria *Nasuia* (in leafhoppers and treehoppers), *Zinderia* (in spittlebugs), *Vidania* (in planthoppers), and alphaproteobacterium *Hodgkinia* (in cicadas) (McCutcheon et al., 2009; McCutcheon and Moran, 2010; Urban and Cryan, 2012). A long time ago, these symbionts reached a relatively stable evolutionary phase, characterized by limited changes in organization, or additional losses of function, despite high rates of nucleotide sequence evolution (McCutcheon et al., 2019; Moran et al., 2008). However, these symbioses are far from stable, with the relatively frequent acquisition of new microbes and symbiont replacement driving additional changes (Sudakaran et al., 2017). Well-known examples of the replacement of an ancient symbiont include *Sodalis* instead of *Zinderia* in Philaenini spittlebugs (Koga et al., 2013); *Baumannia* that accompanies *Sulcia* in sharpshooter leafhoppers (Bennett et al., 2014); or repeated replacements of *Hodgkinia* by *Ophiocordyceps* fungi in various lineages of cicadas (Matsuura et al., 2018). Additional co-infecting microbes, especially *Gammaproteobacteria* related to *Sodalis* and *Arsenophonus*, have also been reported from many Auchenorrhyncha lineages and are typically linked to nutrition - although evidence is often lacking, and the nature and stability of these associations unclear (Kobiałka et al., 2016; Michalik et al., 2021, 2014). Besides these nutritional symbionts, many sap-feeding insects also harbor facultative ones, which are not necessary for survival but can affect many aspects of host biology (Oliver et al., 2010; Zytynska et al., 2021).

Planthoppers (Fulgoromorpha) are an ecologically and evolutionarily diverse Auchenorrhyncha group encompassing almost 14000 known species representing 21 extant families (Bourgoin, 2022). They represent various degrees of trophic specificity and ecological relationships. Due to their food preferences and modes of feeding, planthoppers are vectors of plant pathogens, and several are considered serious agricultural pests (Wilson and O’Brien, 1987). We know that these insects have been ancestrally associated with bacterial symbionts *Sulcia* and *Vidania* (Bennett and Mao, 2018; Michalik et al., 2021; Urban and Cryan, 2012). However, in many planthopper lineages, they are now accompanied or have been replaced by other heritable bacteria (e.g., *Sodalis, Arsenophonus, Purcelliella, Wolbachia*) or fungi (Bennett and Mao, 2018; Michalik et al., 2009, 2021). These associations and modes of their transmission can be very diverse, prompting Paul Buchner, after decades of observations, to famously describe them as “a veritable fairyland of symbiosis” (Buchner, 1965). However, our understanding of the diversity and evolution of these associations is grossly incomplete. Before the Second World War, Hans Müller and Paul Buchner have characterized many planthopper-symbiont associations using histological techniques and interpreted the patterns with impressive accuracy, but lacked tools to verify their identity (Buchner, 1965; Müller, 1940a, 1940b). More recently, sequencing-based approaches have provided information about the identity of some of these microbes (Bennett and Mao, 2018; Michalik et al., 2021), but we know much less about their genomics characteristics, evolutionary patterns, or functions. Importantly, microscopy and sequencing surveys have not generally been combined, and the use of modern microscopy techniques is limited to a few taxa (Michalik et al., 2021). Hence, we lack the link between the host and symbiont identity, the organization of symbiont-holding tissue, and symbiosis functions.

We know that ancient, obligate symbionts generally inhabit bacteriomes, dedicated organs made up of specialized insect cells termed bacteriocytes (Buchner, 1965). As bacteriocytes repeatedly evolved in many insect lineages, bacteriome organization varies between host clades, and often also among different symbionts that live in the same host (Buchner, 1965). The newly-acquired microorganisms can live both within bacteriomes and other tissues. The compartmentalization of symbionts into host cells allows them, on the one hand, to shelter from the insect’s immune system cells; on the other hand, it ensures the host control of the symbiont population growth (Harris et al., 2010). In multi-symbiotic systems, the spatial arrangement of obligate, nutritional symbionts may also reflect the metabolic convergence between them (Douglas, 2016). Additionally, as symbioses evolve and ancient symbiotic associates get complemented or replaced by others, the organization of the symbiont-containing tissue is likely to change further. However, little is known about the localization of newly-acquired microorganisms or how it is determined, even though it significantly influences the outcome of the symbiotic interaction.

Here, we aimed to fill this knowledge gap by exploring the symbiotic systems of 44 planthoppers species of 15 families, representing the main evolutionary lineages of modern Fulgoromorpha. We combined light, fluorescence, and transmission electron microscopy with phylogenomic data to compare the bacteriomes’ organization across planthoppers hosting different symbiont combinations. This approach allowed us to identify and describe seven different categories of symbiont localization within the host insect’s body and seven types of bacteriome organization. We emphasized the localization of microorganisms complementing the ancestral symbionts and newly-acquired microorganisms that replaced the ancestral associates.

## 2. Methods

### 2.1. Insect collection

Taxonomic sampling included 44 planthopper species representing 15 families, including Acanalonidae (1), Achilidae (2), Caliscelidae (2), Cixiidae (3), Delphacidae (7), Derbidae (3), Dictyopharidae (7), Fulgoridae (3), Flatidae (2), Issidae (6), Lophopidae (1), Meenoplidae (1), Ricanidae (3), Tettigometridae (2), and Tropiduchidae (1). Adult insects were collected in Bulgaria, Italy, Vietnam, and Poland between 2014 and 2019. After sampling, the material was preserved in an appropriate fixative (ethanol or glutaraldehyde) and stored at 4°C until further processing. Sampling details are summarized in Supplementary Table S1. Representative specimens from each species were identified based on morphological characteristics.

### 2.2. Sequencing-based symbiont survey

#### 2.2.1. DNA extraction and metagenomic library preparation and sequencing

DNA was extracted from dissected bacteriomes or insect abdomens using one of three different DNA extraction kits: Sherlock AX isolation kit (A&A Biotechnology, Poland), Bio-Trace DNA Purification Kit (Eurx, Poland), and Genomic Mini AX Yeast Spin kit (A&A Biotechnology, Poland), according to manufacturers” protocols. The metagenomic libraries for high-throughput sequencing on the Illumina platform were prepared using NEBNext Ultra II FS DNA Library Prep and NEBNext DNA Ultra II kits, with a target insert length of 350 bp. The pooled libraries were sequenced on Illumina HiSeq X or NovaSeq 6000 S4 lanes (2×150bp reads).

#### 2.2.2. Metagenome-based reconstruction of microbiome composition

The taxonomic composition of the microbial symbiont community was assessed based on the sequences of small subunit rRNA genes. We reconstructed rRNA sequences using PhyloFlash v3.4 (Gruber-Vodicka et al., 2020), using the option --everything, including EMIRGE approach (Miller et al., 2010). The taxonomic classification was performed against a customized SILVA database (v1.38), which included several so-far-unpublished sequences of planthopper symbionts. The results were manually verified through comparisons of sequences and symbiont identities among related species and microscopy and marker gene-sequencing datasets for the additional specimens (data not shown).

#### 2.2.3. Host mitogenome assembly and phylogenetic analysis

The metagenomic reads were assembled using MEGAHIT v1.1.3 (Li et al., 2016) with k-mer size from 99 to 255. The contigs of host mitochondrion were identified using “blastn” and” blastx” searches against custom databases, which included DNA and amino acid sequences of published planthopper mitogenomes. The identified mitochondrial contigs were then annotated with a custom Python script modified from (Łukasik et al., 2019). The script first extracted all the Open Reading Frames (ORFs) and their amino acid sequences from a genome. Then, the ORFs were searched recursively using HMMER v3.3.1 (Eddy, 2011), through custom databases containing manually curated sets of protein-coding and rRNA genes of planthopper mitochondria. rRNA genes were searched with nhmmer (HMMER V3.3.1) (Wheeler and Eddy, 2013), and tRNAs were identified with tRNAscan-SE v2.0.7 (Chan et al., 2021). For phylogenetic reconstructions, we used concatenated alignments of 13 mitochondrial protein coding genes (*nad2, cox1, cox2, atp8, atp6, cox3, nad3, nad5, nad4, nad4L, nad6, cob*, and *nad1*) and one mitochondrial ribosomal RNA (*rrnL*), resulting in a total dataset length of 11713 bp.

The maximum likelihood tree of host species was constructed in IQ-Tree on XSEDE (Minh et al., 2020) and implemented in CIPRES v.3.3 (Miller et al., 2010). ‘Model Selection’ (Kalyaanamoorthy et al., 2017) was selected to search for the best model in CIPRES. The partition type was set to allow the 14 partitions (one for each marker) to allow different evolutionary rates (Chernomor et al., 2016). ‘TESTNEWMERGE’ was specified to allow partitions with similar speed to be analyzed as a single partition. The best fit models were decided by the highest BIC (Bayesian Information Criterion) scores. Bootstrapping was conducted using ‘SH-aLRT’ bootstrap methods with 1000 replicates. All other options were set to default.

#### 2.2.4. Amplicon-based reconstruction of intraspecific microbiome diversity

For selected species that were represented by multiple individuals in our collection, we sequenced amplicons for the V4 region of the 16S rRNA gene as a means of assessing intra-species microbiome diversity alongside the host mitochondrial cytochrome oxidase I (COI) gene as a means of confirming host identity. We described the laboratory workflows previously (Michalik et al. 2021). We used a two-step PCR library preparation protocol, where in the first round of PCR, we simultaneously amplified marker regions of interest using template-specific primers 515F/806R and COIBF3/COIBR2 with Illumina adapter stubs, and then used bead-purified PCR products as the template for the second, indexing PCR. Pooled libraries were sequenced on an Illumina MiSeq v3 lane (2×300-bp reads) at the Institute of Environmental Sciences of Jagiellonian University. We processed amplicon data separately for both targeted regions using custom pipeline based on USEARCH/VSEARCH. Reads assembled into contigs were quality-filtered, then dereplicated and denoised, aligned against the reference databases, screened for chimeras using UCHIME, classified taxonomically, and finally, clustered at 97% identity level using the nearest-neighbor algorithm and divided into OTUs. The COI data were used to confirm insect species identity, while data on relative abundance of symbiont types was visualized using R v. 4.0.2 (R Development Core Team) with the ggplot2 package (Wickham, 2011).

### 2.3. Microscopic analyses

#### 2.3.1. Histological and ultrastructural analyses

Adult specimens of each species were dissected partially in the field and immediately preserved in 2.5% glutaraldehyde in 0.1 M phosphate buffer. In the laboratory, the fixed material was rinsed three times in the same phosphate buffer with the addition of sucrose (5.8g /100ml) and then postfixed in 1% osmium tetroxide. Next, samples were dehydrated in the graded series of ethanol (30-100%) and acetone and embedded in epoxy resin Epon 812 (SERVA, Heidelberg, Germany). Semithin sections were stained with 1% methylene blue in 1% borax and examined using the Nikon Eclipse 80i light microscope. Ultrathin sections were contrasted with uranyl acetate and lead citrate and analyzed using JEOL JEM 2100 electron transmission microscope.

#### 2.3.2. Fluorescence *in situ* hybridization

The Fluorescence *in situ* hybridization was performed with symbiont-specific probes complementary to their 16S rRNA gene sequences (see Table S2). Insects preserved in ethanol were rehydrated and then postfixed in 4% paraformaldehyde for two hours at room temperature. Next, the material was dehydrated again by incubation in increased concentrations of ethanol (30-100%) and acetone, embedded in Technovit 8100 resin (Kulzer, Wehrheim, Germany), and cut into semithin sections. The sections were then incubated overnight at room temperature in a hybridization buffer containing the specific sets of probes. After hybridization, the slides were washed in PBS three times, dried, covered with ProLong Gold Antifade Reagent (Life Technologies), and examined using a confocal laser scanning microscope Zeiss Axio Observer LSM 710.

## 3. Results

### 3.1. High-throughput sequencing reveals the diversity of planthopper symbioses

The reconstruction of microbial community composition based on metagenome-derived, full-length bacterial 16S rRNA and fungal 18S rRNA sequences revealed a striking variety of symbioses across 44 sampled species from 15 planthopper families (Fig. 1). Specifically, Phyloflash recovered 16S rRNA sequences of 15 bacterial genera representing four phyla: *Bacteroidetes* (*Sulcia, Cardinium*), *Proteobacteria* (*Vidania, Purcelliella, Sodalis, Arsenophonus, Wolbachia, Rickettsia, Pectobacterium, Asaia*-like symbionts, *Serratia, Sphingomonas*, and *Pantoea*), *Actinobacteria* (*Frigoribacterium*), and *Tenericutes* (*Spiroplasma*). It also reconstructed sequences of fungi in the order Hypocreales (phylum Ascomycota). Generally, symbiont community composition in Fulgoromorpha is linked to host phylogeny, but with some exceptions, including families Achilidae and Issidae.

**Figure 1.**
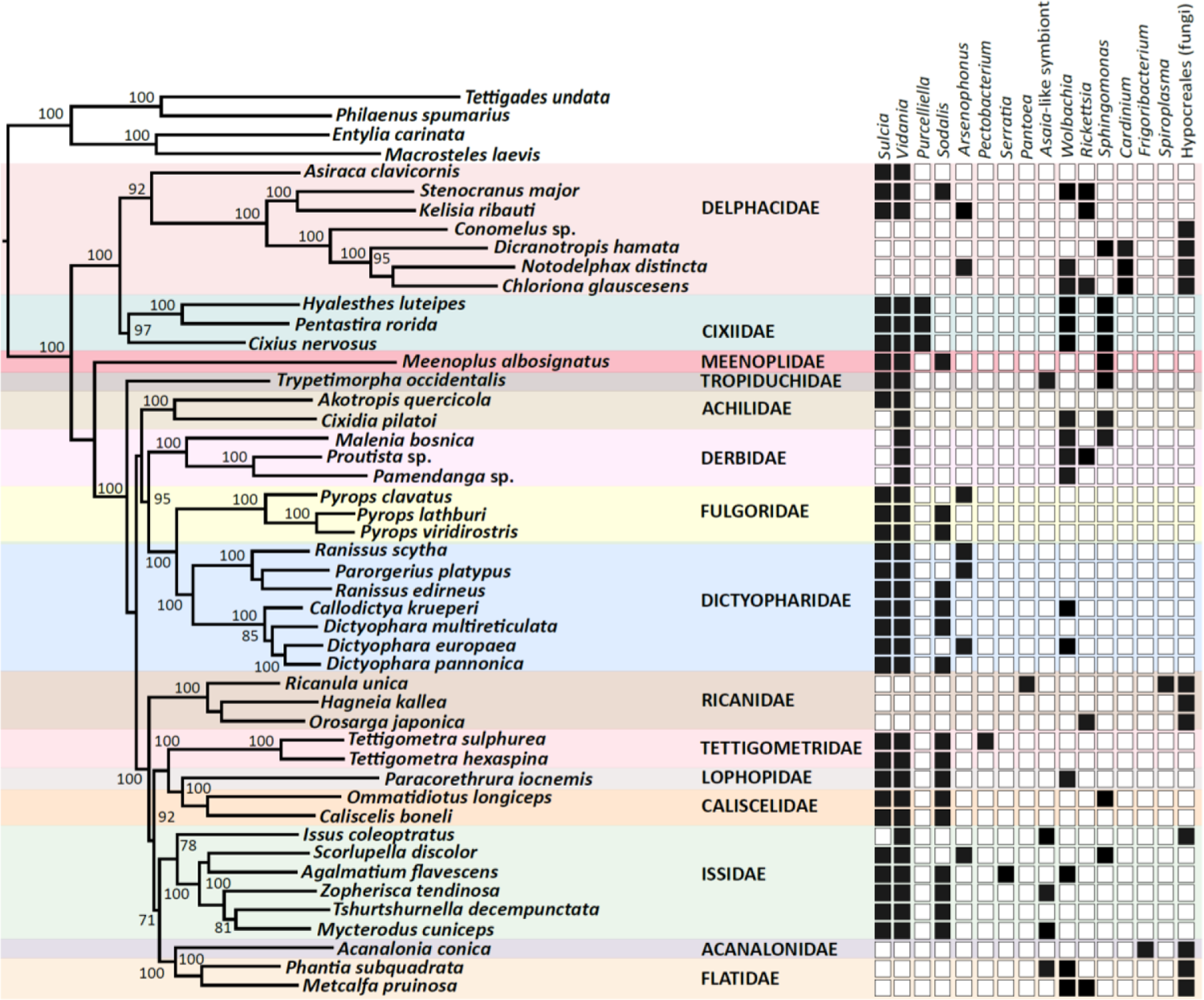
Symbiont diversity across planthopper phylogeny. The Maximum Likelihood phylogeny is constructed based on concatenated 13 mitochondrial protein-coding genes and one mitochondrial ribosomal RNA, of the total size of 11713 bp. Bootstrap support values >70% are shown. The heatmap shows the presence/absence planthopper symbiont genera, based on the sequences of 16S rDNA genes reconstructed from metagenomic datasets.

The most broadly distributed bacterial symbiont in Fulgoromorpha is the ancient betaproteobacterial nutritional endosymbiont *Vidania*, detected in 34 species representing 12 families. In 9 of these families, including Caliscelidae, Cixiidae, Delphacidae, Dictyopharidae, Fulgoridae, Lophopidae, Meenoplidae, Tettigometridae, and Tropiduchidae, *Vidania* always co-resides with *Sulcia*. In contrast, in the families Achilidae and Issidae, some species host both these ancestral symbionts while others only harbor *Vidania*. Derbidae is the only family where all tested representatives lack *Sulcia* but host ancestral *Vidania*. In all examined species of families Acanalonidae, Flatidae, Ricanidae, and some members of the family Delphacidae (subfamily Delphacinae), we did not observe *Sulcia* or *Vidania*, and detected fungal symbionts belonging to the order Hypocreales instead (Fig. 1).

Planthoppers hosting ancestral symbionts *Sulcia* and *Vidania* are associated with at least one additional bacterium, which in most species belong to the class *Gammaproteobacteria*. Among them, *Sodalis* and *Arsenophonus* are the most common, colonizing 17 and 7 species in 8 and 4 families, respectively. Another gamma-symbiont, *Purcelliella*, occurs exclusively in members of the family Cixiidae. Other gammaproteobacterial associates found in some examined planthoppers include *Pectobacterium, Serratia*, and *Pantoea*. Fulgoromorpha are also frequently colonized by microbes belonging to the class *Alphaproteobacteria. Wolbachia* and *Rickettsia*, known as facultative endosymbionts of diverse insects, are together detected in 8 out of 15 families. In 5 species: *Issus coleoptratus, Mycterodus cuniceps, Phantia subquadrata, Trypetimorpha occidentalis*, and *Zopherisca tendinosa*, we also identified alphaproteobacteria from the family Acetobacteraceae. Bacteria limited to species harboring fungal symbionts represent the genera *Cardinium* (in *Dicranotropis hamata* and *Notodelphax distincta*), *Frigoribacterium* (in *Acanalonia conica*), and *Spiroplasma* (in *Ricanula unica*) (Fig. 1).

In some of 30 planthopper species where multiple individuals per population were used for amplicon sequencing, we also observed intra-species diversity in the microbiome composition (Fig. 2). *Sulcia, Vidania*, and fungal symbionts were uniformly either present or absent in all individuals of a species. In contrast, in some species, we observed differences in the composition of additional symbionts (e.g., *Wolbachia, Rickettsia*, and *Sphingomonas*) and their relative abundance between individuals sampled from one population. For example, *Asaia*-like symbiont was detected in only 2 out of 3 individuals of *Issus coleoptratus*, while the third hosted *Sphingomonas* but lacked *Asaia*.

**Figure 2.**
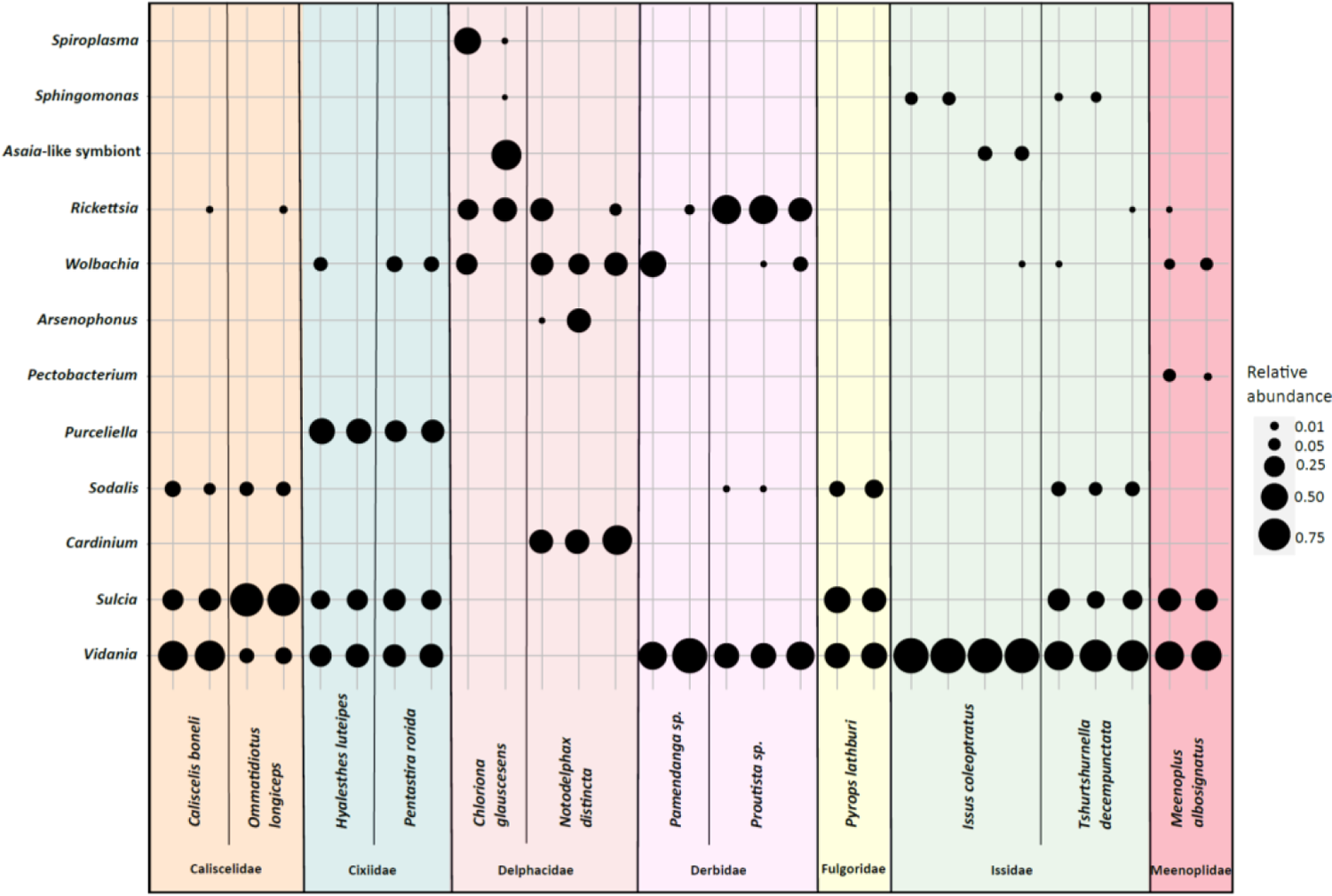
The diversity of symbionts in replicate individuals of selected planthopper species. The blob sizes correspond to the relative abundance of a symbiont from a given bacterial genus.

### 3.2. Microorganisms associated with planthoppers show different localization patterns

Our broad microscopic investigations have revealed that symbionts associated with planthoppers are distributed in very different ways across host insect tissues (Fig. 3). Most planthopper symbionts reside in specialized organs (bacteriomes or mycetomes), which can differ considerably in their organization depending on microorganisms hosted (Fig. 4). Below, we present different symbiont tissue localizations and then describe different bacteriome/mycetome types. In subsequent sections, we explain how different symbiont taxa are distributed across these localizations and symbiont-containing organ types in different planthopper clades.

**Figure 3.**
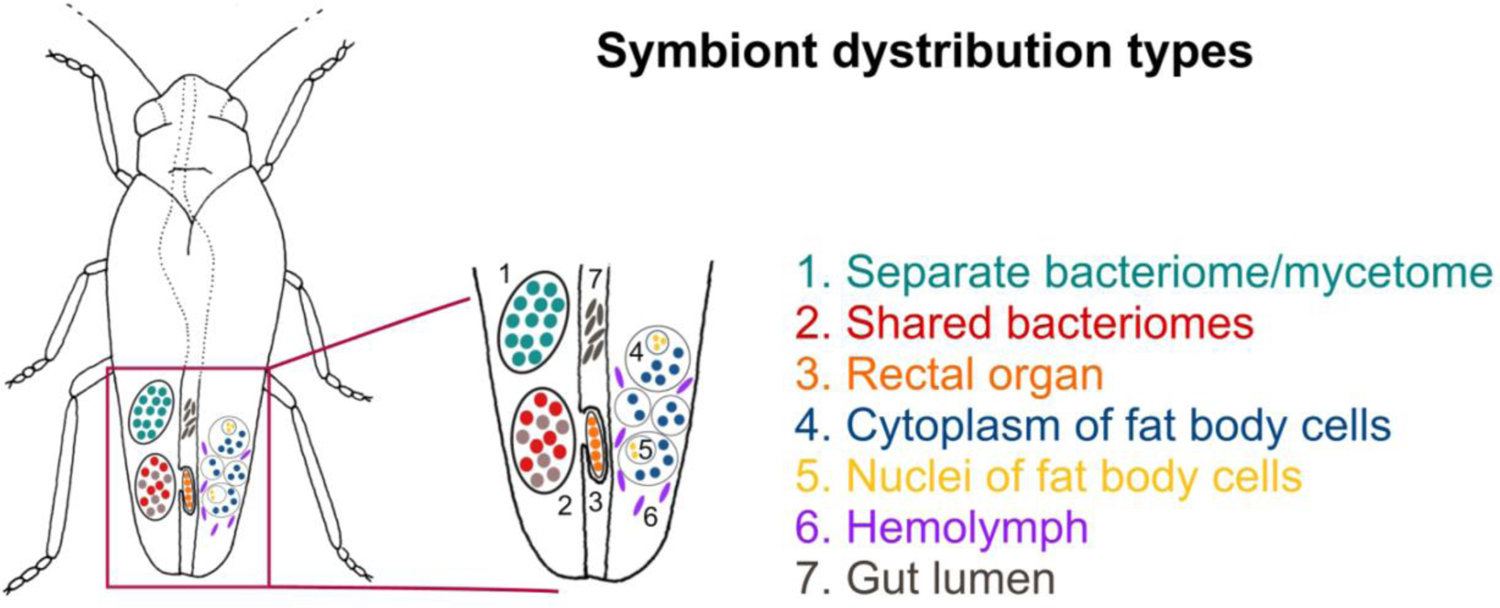
Schematic representation showing the possible symbiont localizations in host-insect body.

**Figure 4.**
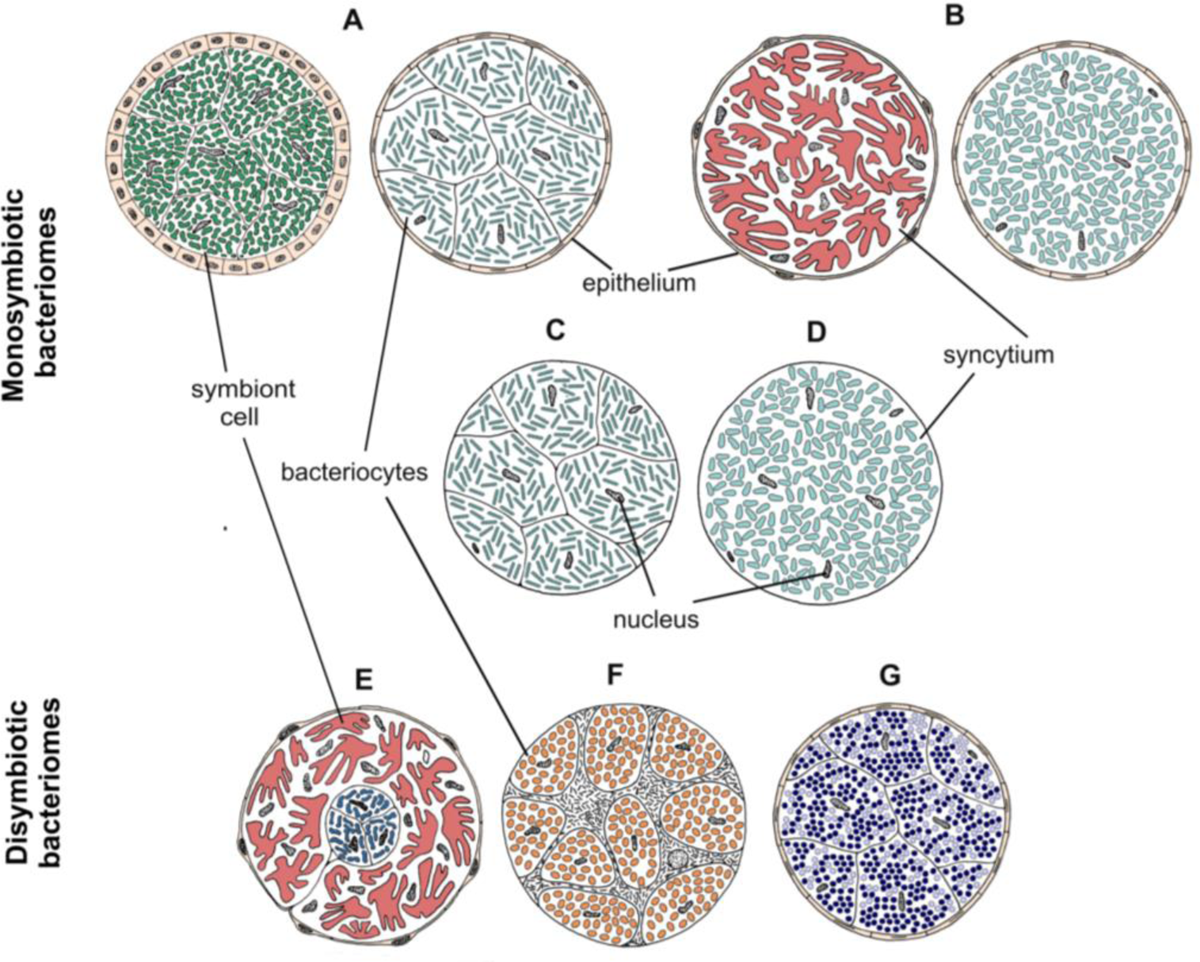
Drawings showing the organization of different types of bacteriomes and mycetomes in planthoppers.

We identified seven distinct localizations of symbionts (Fig. 3). Generally, (1) different symbionts are segregated to separate bacteriomes; however, in some cases, (2) they share a common bacteriome. In females only, in addition to the the bacteriomes localized in the body cavity, (3) a single bacteriome called rectal organ occurs in the deep invagination of the hindgut. Planthopper symbionts may also be distributed in the fat tissue, where they may reside in (4) the cytoplasms of the fat body cells or (5) fat body cell nuclei. Besides intracellular localization, microorganisms associated with planthoppers may occur extracellularly: either (6) in the hemolymph between fat body cells or (7) in the gut lumen.

The most common localization of planthopper symbionts are bacteriomes - structures usually consisting of bacteriocytes filled with symbionts. Analogous organs filled with fungal cells are known as mycetocymes/mycetocytes, but as their organization does not depart from that of bacteriomes, we will only write about bacteriocytes for simplicity. Basically, bacteriomes are large, elongated, paired or unpaired structures that are localized in the insect’s body cavity in the abdomen. In males, bacteriomes are much smaller than in females and are always localized in the rearmost portion of the abdomen (not shown). In contrast, in females, bacteriomes are distributed close to the ovaries and show different spatial arrangements. Usually, they run longitudinally or transversely through the posterior part of the abdomen, but sometimes they are intertwined with the ovaries (not shown). Based on the comparative analysis of bacteriome organization, we distinguished 7 types of bacteriomes in planthoppers, differing in structure and number of symbionts inhabiting them (Fig. 4). The first four types (A-D) refer to bacteriomes containing only one symbiont type. Type A bacteriome comprises several mononucleated bacteriocytes and is surrounded by a single layer of epithelial cells. The thickness of epithelium varies among bacteriomes harboring different symbionts. Type B represents plasmodium-like, nuclei-rich syncytial bacteriomes covered by the epithelium of varying thickness (similar to type A). Bacteriome types C and D are not surrounded by epithelial bacteriome sheath. Type C is made up of several bacteriocytes, whereas type D is a multi-nuclei syncytium.

The remaining three types (E-G) refer to bacteriomes harboring two different microorganisms, generally with more complex organization. Bacteriome type E comprises several closely adjacent bacteriocytes filled with symbiont 1, which are covered by a single syncytial bacteriome harboring symbiont 2. In turn, bacteriome type F shows a unique structure that can be described as “cells enclosed in a cell”. The bacteriome is a big multi-nuclei cell filled with cells of symbiont 1 and bacteriocytes containing symbiont 2. In the last type of bacteriome (G), two kinds of symbionts are mixed in the cytoplasms of one bacteriocyte.

### 3.3. Ancestral symbionts *Sulcia* and *Vidania* retain conserved morphology and localization

The two ancestral symbionts of planthoppers, *Sulcia* and *Vidania*, are restricted to bacteriomes in all examined planthopper species. These two symbionts always reside in separate bacteriomes (Fig. 5). Bacteriomes occupied by *Sulcia* represent type A. They are paired, tubular, and covered by a thick monolayered epithelium termed bacteriome sheath (Fig. 5A-C). The epithelial cells are cube-shaped and have large, spherical nuclei and numerous mitochondria in the cytoplasm (Fig. 5A, C). The bacteriocytes that make up the bacteriome are uninucleated and closely adhere to each other. Their cytoplasm is filled with variably shaped (pleomorphic) cells of *Sulcia* (Fig. 5A-C). Basically, the *Sulcia* cell shape and size are similar among planthoppers, with one exception for *Kelisia* and *Stenocranus* genera in which *Sulcia* cells are smaller and more spherical (not shown).

**Figure 5.**
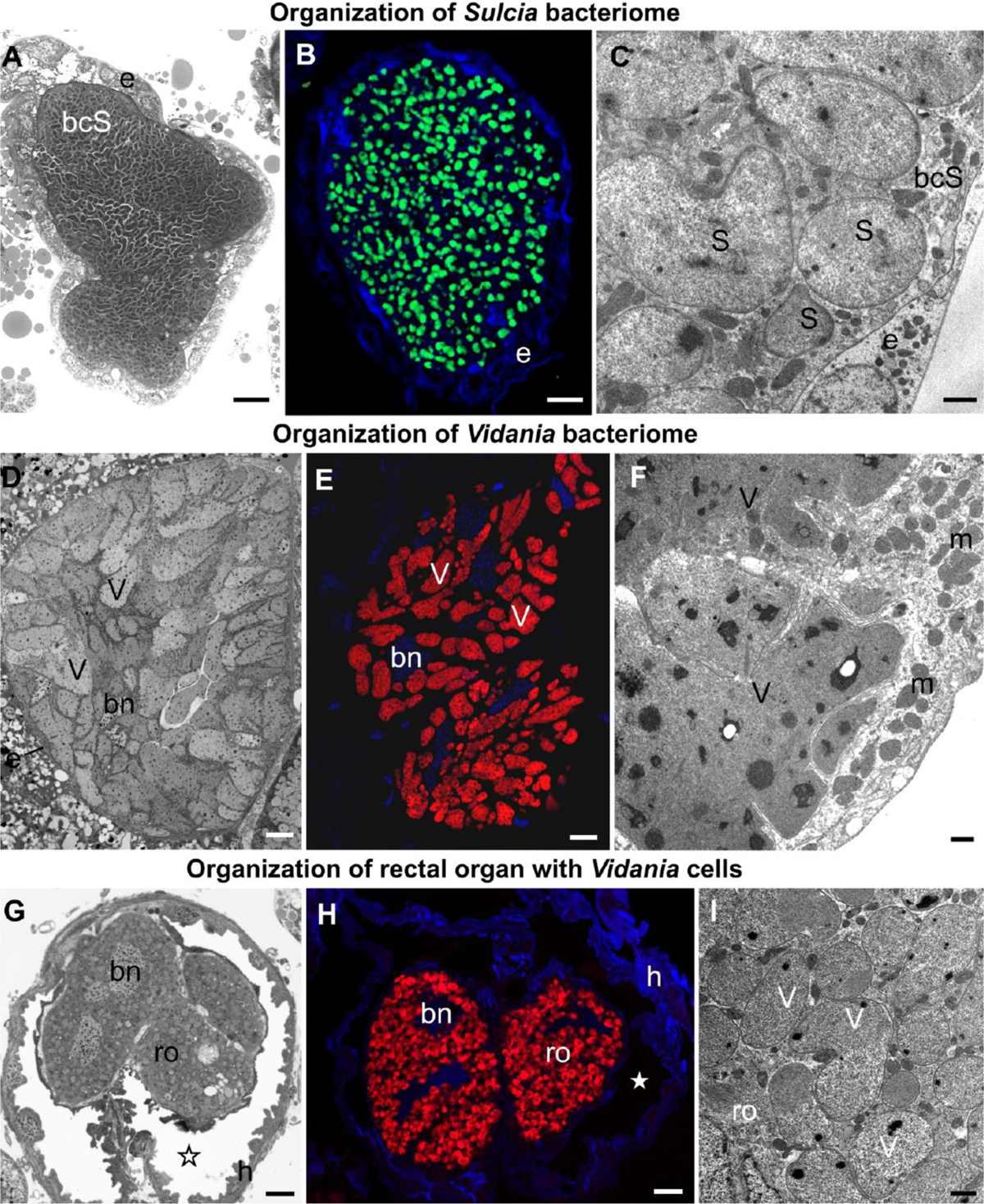
Bacteriome structure and morphology of ancient planthopper symbionts *Sulcia* and *Vidania*. **A**. Organization of *Sulcia* bacteriome **B**. Visualisation of bacteriomes inhabited by *Sulcia* using *Sulcia*-specific probe (green), blue represents DAPI, **C**. Ultrastructure of *Sulcia* cells. **D**. Organization of the *Vidania* bacteriome. **E**. Visualisation of bacteriome inhabited by *Vidania* using *Vidania*-specific probe (red), blue represents DAPI **F**. Ultrastructure of *Vidania* bacteriome **G**. Rectal organ with *Vidania* symbiont in the hindgut lumen. **H**. FISH detection of *Vidania* in the rectal organ. **I**. Ultrastructure of *Vidania* cells in the rectal organ. **A. D. G**. Light microscope (LM), scale bar = 10 μm. **B. E. H**. Confocal microscope, scale bar = 10 μm. **C. F. I**. Transmission electron microscope (TEM), scale bar = 1 μm. bcS - *Sulcia* bacteriocyte, bn - bacteriocyte nucleus, e - epithelium, h - hindgut, m - mitochondria, ro - rectal organ, asterisk - gut lumen, S - *Sulcia*, V - *Vidania*. **Insect species: A**. *Tettigometra sulphurea* (Tettigometridae) **B**. *Akotropis quercicola* (Achilidae) **C**. *Meenoplus albosignatus* (Meenoplidae) **D. E**. *Tettigometra sulphurea* (Tettigometridae) **F**. *Ommatidiotus longiceps* (Caliscelidae) **G**. *Proutista* sp. (Derbidae) **H**. *Akotropis quercicola* (Achilidae) **I**. *Cixidia pilatoi* (Cixiidae)

The *Vidania* symbiont is also strictly limited to bacteriomes (Fig. 5D-I). In both sexes, bacteriomes with *Vidania* occur in the body cavity between internal organs. These bacteriomes represent type B, and are large, multi-nucleated syncytial organs surrounded by a very thin, flattened bacteriome sheath (Fig. 5D, F). Their cytoplasm is tightly packed with giant, lobed *Vidania* cells (Fig. 5D-F). The nuclei are usually scattered between *Vidania* cells, whereas the numerous mitochondria mostly lay in the peripheral part of the bacteriome under the bacteriome membrane (Fig. 5E). In addition to these syncytial bacteriomes present in both sexes, females have an additional, unpaired bacteriome called a rectal organ, which is situated in the deep invagination of the hindgut that protrudes into its lumen. The rectal organ is composed of several binucleated bacteriocytes. The *Vidania* cells that fill them are pleomorphic, in sharp contrast to the cells in the primary bacteriome (Fig. 5G-I).

### 3.4. Gammaproteobacterial symbionts usually reside in bacteriocytes

Gammaproteobacterial symbionts, including *Sodalis, Arsenophonus, Purcelliella*, and *Pectobacterium*, usually inhabit bacteriomes. However, the organization of these organs varies among symbiont genera and to some extent, also host clades (Fig. 6).

**Figure 6.**
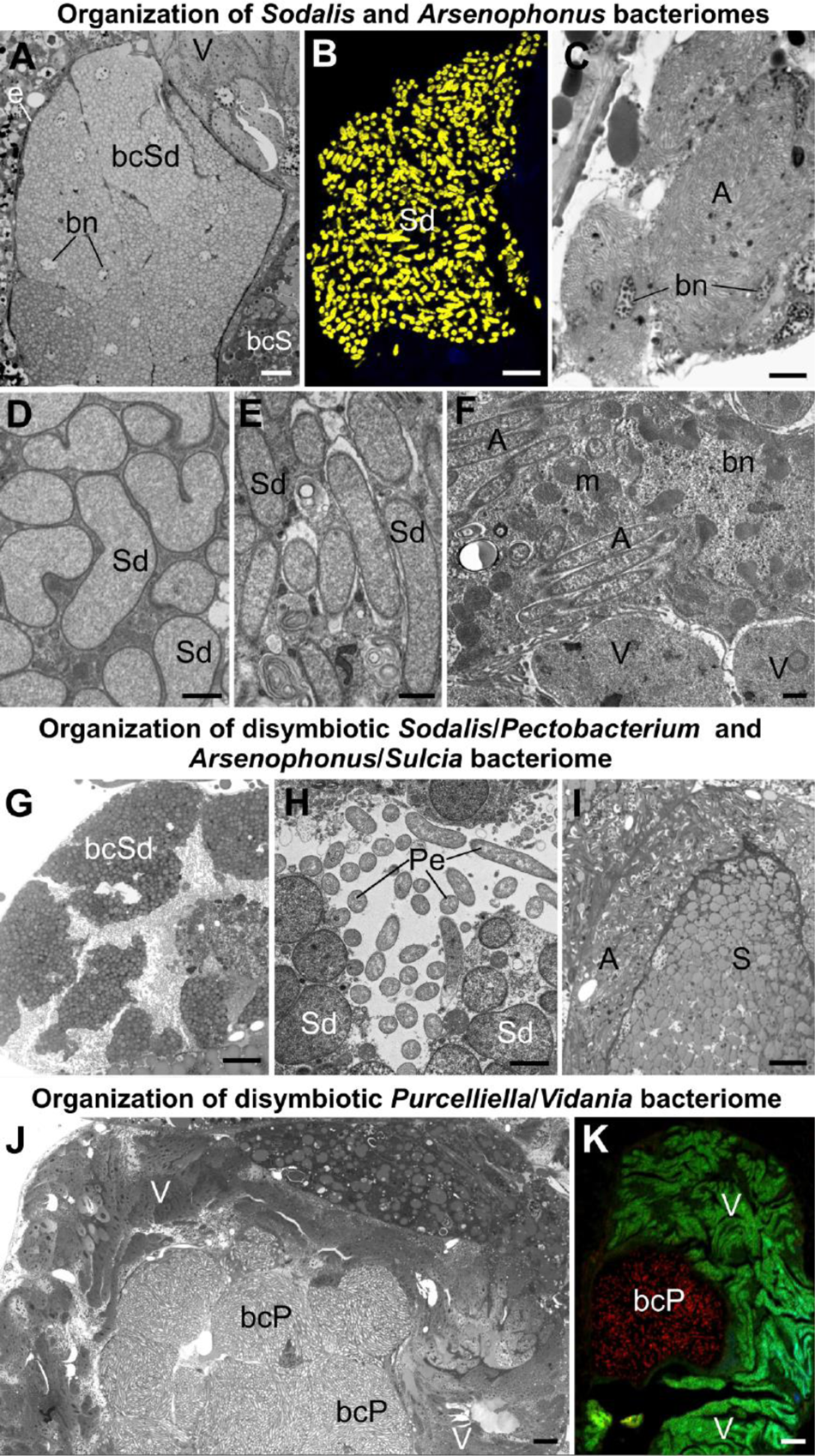
Organization of bacteriomes harboring gammaproteobacterial symbionts in planthoppers. **A. B**. Organization of *Sodalis* bacteriome **C**. Cluster of *Arsenophonus* bacteriocytes, **D. E**. Ultrastructure of *Sodalis* cells **F**. *Arsenophonus* cells in the cytoplasm of *Vidania* bacteriocyte. **G. H**. Organization of disymbiotic *Sodalis*/*Pectobacterium* bacteriome. **I**. Organization of disymbiotic *Arsenophonus*/*Sulcia* bacteriome. **J. K**. Organization of disymbiotic *Purcelliella*/*Vidania* bacteriome. **A. C. G. H. I. K**. Light microscope (LM), scale bar = 10 μm. **B. H**. Confocal microscope, scale bar = 10 μm. **D. E. F. J**. Transmission electron microscope (TEM), scale bar = 1 μm. A - *Arsenophonus*, bcP - *Purcelliella* bacteriocyte, bcSd - *Sodalis* bacteriocyte, bn - bacteriocyte nucleus, m - mitochondria, Pe - *Pectobacterium*, S - *Sulcia*, Sd - *Sodalis*, V – *Vidania* **Insect species: A**. *Tshurtshurnella decempunctata* (Issidae) **B**. *Tettigometra griseola* (Tettigometridae) **C**. *Scorupella discolor* (Issidae) **D**. *Zopherisca tendinosa* (Issiade), **E**. *Kelisia ribauti* (Delphacidae) **F**. *Scorupella discolor* (Issidae) **G. H**. *Tettigometra sulphurea* (Tettigometridae) **I**. *Pyrops clavatus* (Fulgoridae) **J. K**. *Hyalesthes luteipes* (Cixiidae)

The most common planthopper gammaproteobacterial symbionts, *Sodalis* and *Arsenophonus*, in almost all cases, occur in separate bacteriomes (Fig. 6A-C). These bacteriomes are unpaired, composed of closely adhering bacteriocytes (types C and D), and are or are not covered by epithelial sheath. The bacteriocytes are usually binucleated and tightly packed with bacterial cells (Fig. 6A, B, E-F). The exceptions from this general rule were observed in *Scorupella discolor* (Issidae), and *Pyrops clavatus* (Fulgoridae). In *S. discolor* bacteriocytes harboring *Arsenophonus* symbionts are not integrated into compact bacteriome but form a more or less loose cluster of cells (Fig. 6C). In this species *Arsenophonus* symbionts may also reside in the cytoplasm of *Vidania’s* bacteriocytes, with both symbionts mixed in the cytoplasm of the shared bacteriocytes (type G) (Fig. 6F). In turn, in *P. clavatus Arsenophonus* symbionts surround the *Sulcia* bacteriome (type E) (Fig. 6I).

Apart from the differences in bacteriome organization, we also observed differences in *Sodalis* cell shape - from rod-shaped (in Dictyopharidae) through irregular (in Issidae) to almost spherical (in Tettigometridae) (Fig. 6A, B, D, E, G, H). Furthermore, in the cytoplasm of bacteriocytes with *Sodalis* in different planthoppers species, we observed numerous lamellar bodies, which we interpret as symptoms of *Sodalis* degeneration (Fig. 6E).

Other gammaproteobacterial symbionts - *Purcelliella* and *Pectobacterium*, share the bacteriome with another symbiont. *Purcelliella* - a symbiont exclusive to the family Cixiidae, occurs in a common bacteriome with *Vidania* symbiont (type E). *Purcelliella* inhabits separate bacteriocytes, but they are always covered by the large syncytial bacteriome with *Vidania* cells (Fig. 6J, K). In turn, in *Tettigometra sulphurea, Pectobacterium* and *Sodalis* co-reside in the common bacteriome with the most complicated organization we observed in planthoppers (type F). This bacteriome is a large multinucleate cell cell; within its cytoplasm we observed *Pectobacterium* cells, as well as bacteriocytes with *Sodalis* (Fig. 6G, H).

### 3.5. Alphaproteobacterial symbionts may occupy diverse tissue and organs

Alphaproteobacterial symbionts of planthoppers include the bacteria *Rickettsia, Wolbachia, Sphingomonas*, and bacteria related to *Asaia*. Their localization is not restricted to the bacteriomes - they may occur in other insects’ organs and tissue. Besides bacteriomes, we found alphaproteobacterial symbionts also in the cytoplasm of fat body cells, in the nuclei, gut epithelium, salivary glands, and in females, in different parts of the reproductive system, which is probably related to the symbionts’ transovarial transmission between generations. Alphaproteobacterial symbionts are also the only ones occurring extracellularly in the gut lumen and hemolymph (Fig. 7).

**Figure 7.**
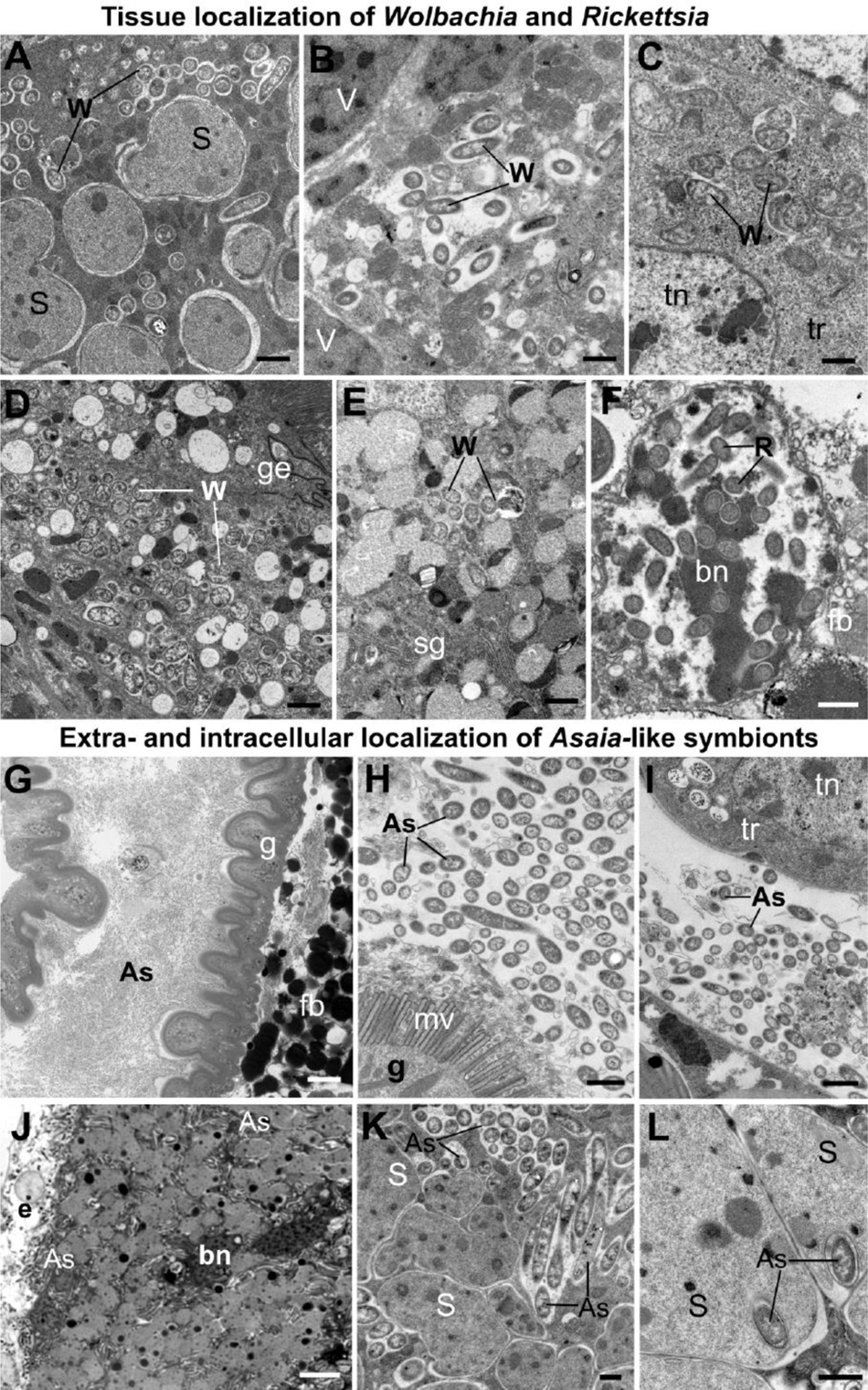
Tissue localization of alphaproteobacterial symbionts in planthoppers. **A-F**. Tissue localization of *Wolbachia* and *Rickettsia* symbionts. **A**. *Wolbachia* mixed with *Sulcia* cells in the shared bacteriome. **B**. *Wolbachia* in the cytoplasm of *Vidania* bacteriocyte. **C**. Wolbachia cells in the tropharium (part of the ovary). **D**. *Wolbachia* in the gut epithelium **E**. *Wolbachia i*n the salivary gland **F**. *Rickettsia* in the nucleus of fat body cell. **G-L**. Extra- and intracellular localization of *Asaia*-like symbionts. **G. H**. *Asaia*-like symbionts in the gut lumen. **I**. *Asaia*-like cells in the hemolymph. **J**. Bacteriome harboring *Sulcia* and *Asaia*-like symbionts **K**. *Sulcia* cells and *Asaia*-like symbionts in the cytoplasm of shared bacteriocyte. **L**. *Asaia*-like cells surrounded by *Sulcia* projections. **A.-F. H. I. L**. Transmission electron microscope (TEM), scale bar = 1 μm. **G. J. K**. Light microscope (LM), scale bar = 10 μm. As - *Asaia*-like symbiont, bn - bacteriocyte nucleus, fb - fat body cell, fn - fat body cell nucleus, g - gut, ge - gut epithelium, mv - gut microvilli, R - *Rickettsia*, S - *Sulcia*, tn - trophocyte nucleus, tr - tropharium, W - *Wolbachia*. **Insect species: A**. *Dictyophara pannonica* (Dictyopharidae) **B**. *Hyalesthes luteipes* (Cixiidae) **D**. *Pentastira rorida* (Cixiidae) **E**. *Ommatidiotus longiceps* (Calliscellidae) **F**. *Orosanga japonica* (Ricanidae) **G. I**. *Phantia subquadrata* (Flatidae) **J-L**. *Trypetimorpha occidentalis* (Tropiduchidae)

The most common alphaproteobacterial symbionts in planthoppers - *Wolbachia* and *Rickettsia*, usually co-occur in the bacteriocytes with *Sulcia* and *Vidania*. The organization of these bacteriomes is similar to the organization of bacteriomes inhabited by *Sulcia* and *Vidania* (type A and B), respectively. The only difference is that in the bacteriocyte cytoplasm, two types of symbionts are mixed (Fig. 7A, B, J, K). Alphaproteobacterial symbionts may also be dispersed in the fat body tissue. Symbionts usually occupy the cytoplasm of fat body cells, but their abundance and density differ between species. *Wolbachia* and *Rickettsia* localized in the cytoplasm of fat body cells are not very numerous and do not occur in all cells. *Rickettsia* associated with the planthopper *Orosanga japonica* (Ricaniidae) has a unique localization: we found it exclusively in the nuclei of fat body cells. In all specimens of that species examined, we observed several *Rickettsia* cells inside the nuclei (Fig. 7F).

Among alphaproteobacterial symbionts detected in planthoppers, *Asaia*-like symbionts show the greatest diversity of tissue localizations across host insect species. In *P. subsquadrata* and *Z. tendinosa*, they inhabit mainly the gut lumen but are also found in hemolymph (Fig. 7G-I). In turn, in *Trypetimorpha occidentalis, Asaia*-like symbionts occur exclusively in the bacteriocytes with *Sulcia* (Fig. 7J-L). Most of its cells are localized in the bacteriocytes’ cytoplasm among *Sulcia* cells (Fig. 7J, K). However, some are almost completely surrounded by *Sulcia* cell projections, making the localization similar to nested symbiosis observed in other Auchenorrhyncha species (Fig. 7L).

### 3.6. Fungal symbionts occur in the mycetomes or fat body cells

All fungal symbionts detected in examined planthoppers species belong to the order Hypocreales (Fig. 1). However, they colonize different tissues across the surveyed insect host species (Fig. 8). Fungal symbionts associated with Flatidae inhabit large organs termed mycetomes. They are usually large, multinuclear syncytia (type B and D) filled with fungal cells and sometimes surrounded by a one-layered epithelium (Fig. 8A, B, G). In contrast, fungal symbionts present in members of families Ricanidae, some Delphacidae, and *Issus coleoptratus* from the family Issidae are not segregated to the mycetomes but occupy fat body cells. They may be scattered across the whole fat tissue (like in all Ricanidae, *Notodelphax distincta* and *Issus coleoptratus*) (Fig. 8 D, E, F, H) or occupy clusters of fat body cells located between the internal organs and against the body wall (Fig. 8C). In *Orosanga japonica* (Ricanidae), fungal symbionts also show intercellular localization in hemolymph between fat body cells (Fig. 8 F, I).

**Figure 8.**
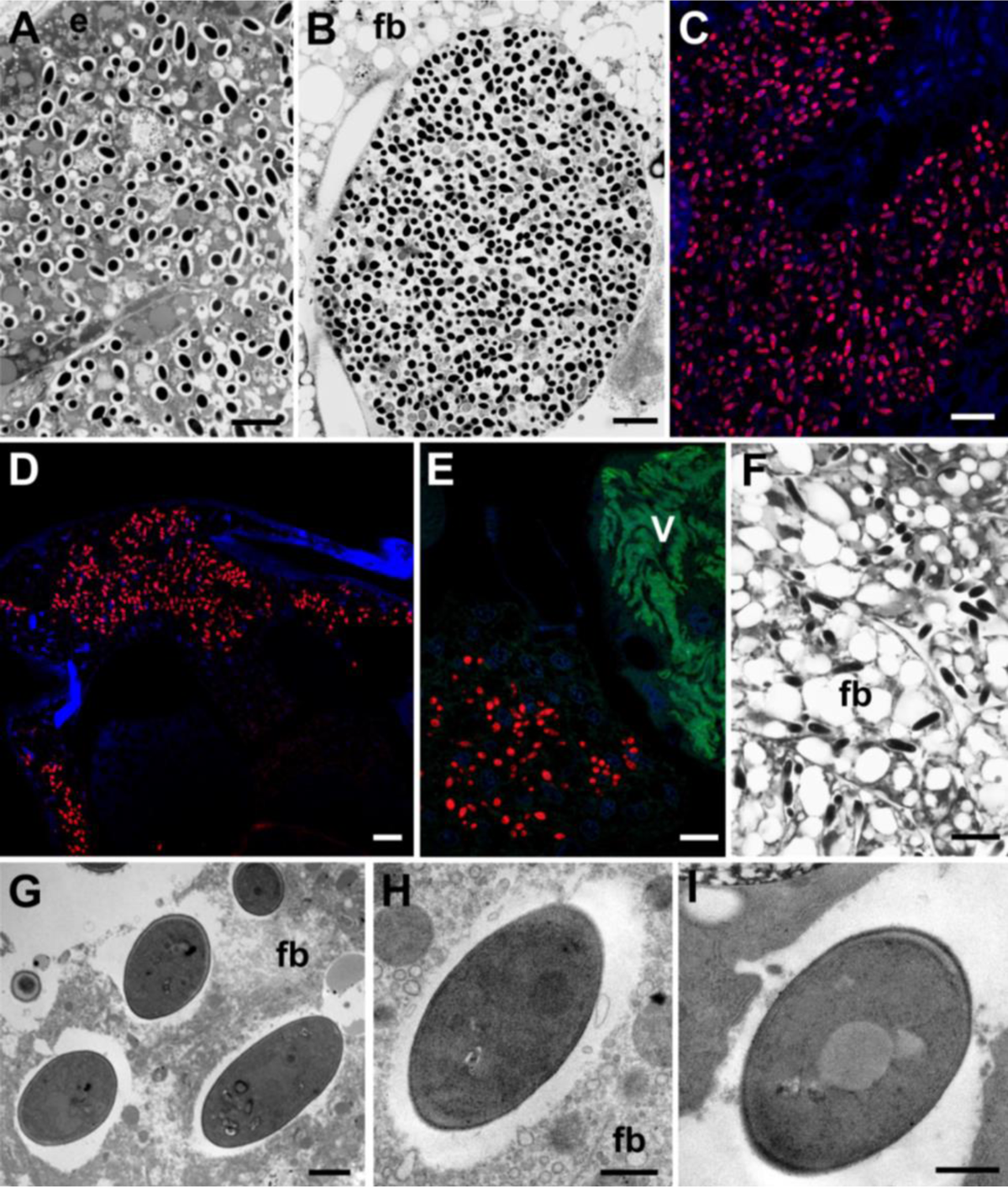
Distribution of fungal symbionts within planthopper tissue. **A-B**. Mycetocyte with fungal symbionts. **C**. Group of fat body cells occupied by fungal symbionts. **D-F**. Fungal symbionts dispersed within fat body cells **G-I**. Ultrastructure of fungal symiont cells. **A.B. F**. Light microscope (LM), scale bar = 10 μm. **C. D. E**. Confocal microscope, scale bar = 10 μm. **G-I**. Transmission electron microscope (TEM), scale bar = 1 μm. e - epithelium, fb - fat body, V – *Vidania* **Insect species: A**. *Metcalpha pruinosa* (Flatidae) **B**. *Phantia subquadrata* (Flatidae) **C**. *Chloriona glauscesens* (Delphacidae) **D**. *Conomelus* sp. (Delphacidae) **E. F**. I*ssus coleoptratus* (Issidae) **G**. *Chloriona glauscesens* (Delphacidae) **H**. *Dicanotropis hamata* (Delphacidae) **I**. *Ricanula unica* (Ricanidae)

All fungal symbionts have similar morphology - their cells are spherical and surrounded by a thick cell wall (Fig. 8G-I).

## 4. Discussion

### 4.1. Symbiont acquisition and replacement as the main driving forces of the planthopper symbionts’ diversity

Essential endosymbiotic associations can be very stable, as evidenced by the long-term conservation of the organization and function of cellular organelles. However, recent research has made it clear that in many organisms, endosymbiosis is an ongoing and dynamic process that strongly influences their biology (Husnik and McCutcheon, 2016; Matsuura et al., 2018; McFadden, 2001; Michalik et al., 2021; Poole and Gribaldo, 2014). Planthoppers serve as a good example of such dynamic processes and patterns.

Like almost all other Auchenorrhyncha, planthoppers rely on the supplementation of their nutritionally imbalanced diet on symbioses dating back some 300 my (Moran et al., 2005; Bennett and Mao, 2018; Michalik et al., 2021). However, the current picture of symbiosis in many Fulgoromorpha departs significantly from the ancestral state.

We showed that the ancient symbionts *Sulcia* and *Vidania* are still present in most planthopper families and a large share of species, detecting *Vidania* in 77% of species examined and *Sulcia* in 66%. These values are higher than Müller (1949) microscopy-based estimates of 58% and 41%, respectively, and Urban and Cryan’s (2012) diagnostic PCR-based estimates (52% and 39%). The discrepancies likely result from different sampling depths across families, combined with sampling different geographic regions, and perhaps methodological biases. Regardless, it is clear that the loss of one or both of the ancestral symbionts took place many times in the evolutionary history of planthoppers and was usually coupled with the acquisition of new microorganisms that took over their biological functions (Fan et al., 2015; Urban and Cryan, 2012).

The replacement of *Sulcia* by *Ophiocordyceps* fungi, observed in *Issus coleoptratus* (Issidae), parallels the observations from some Deltocephalinae and Ledrinae leafhoppers and from cicadas (Kobiałka et al., 2018; Matsuura et al., 2018; Nishino et al., 2016). In cicadas, specialized fungal pathogens replaced the *Hodgkinia* symbiont independently in different clades, taking over its nutritional responsibilities, while *Sulcia* remained in place (Matsuura et al., 2018). However, in planthoppers, fungal replacement of both *Vidania* and *Sulcia* seem to be more common than the replacements of *Sulcia* alone. We found Hypocreales fungi, without *Sulcia* and *Vidania* but typically accompanied by other bacteria, in five planthopper families: Acanalonidae, Flatidae, Ricanidae, and Delphacidae (in subfamily Delphacinae only). Symbiotic systems with fungal symbionts playing the central role was previously documented in Hemiptera other than planthoppers, including the leafhopper *Scaphoideus titanus*, some aphids (Hormaphididae), and scale insects (Coccidae) (Fan et al., 2015; Michalik et al., 2009; Noda et al., 1995; Szklarzewicz et al., 2021).

The loss of *Sulcia* that we observed in families Derbidae and Achilidae, not compensated by the acquisition of other nutrient-providing symbionts, is harder to explain. It seems possible that this monosymbiotic system, unique among Auchenorrhycha (Müller, 1949), is related to the special habitat and food preferences of these planthoppers, thought to feed on fungal hyphae during their larval stage (Howard et al., 2001). Such diet may be more nutritionally balanced than plant sap, eliminating the need to have all essential amino acids provided by the symbionts, and enabling the loss of symbionts that have thus become redundant. The loss of *Sulcia* may be less detrimental for planthoppers than losing *Vidania*, which provides 7 out of 10 essential amino acids (Bennett and Mao, 2018; Deng et al., 2022; Michalik et al., 2021). It is also possible that *Wolbachia*, present in all Derbidae planthoppers that we characterized, contributes to nutrition, as it does in bed bugs and likely, some Dictyopharidae planthoppers (Hosokawa et al., 2010; Michalik et al., 2021).

In fact, microbes other than *Vidania, Sulcia*, and/or fungi, are often likely to contribute to planthopper nutrition. Gammaproteobacterial symbionts are the strongest candidates for important nutritional roles. In all four planthopper symbioses characterized to date using genomics approaches, the genomes of gammaproteobacterial symbionts (*Purcelliella, Sodalis, Arsenophonus*) encoded B vitamin biosynthesis genes (Bennett and Mao, 2018; Michalik et al., 2021) and some amino acid biosynthesis genes. This was despite their independent origins and distinct genomic characteristics, indicative of very different histories of association with hosts. This seems to be a more general trend: some clades of gammaproteobacteria have repeatedly colonized diverse planthoppers, adopting means of transferring vertically, providing deficient nutrients, and replacing symbiont strains that were there before. The genus *Sodalis* is a particularly striking example. It comprises both versatile opportunists capable of infecting humans but encoding an array of biosynthesis genes, as well as heritable endosymbionts of diverse insects that are likely derived from such opportunists (Enomoto et al., 2017; McCutcheon et al., 2019). For instance, Husnik and McCutcheon (2016) proposed that the diversity of genomic characteristics of mealybug endobacterial symbionts inhabiting the cytoplasm of their ancient *Tremblaya* endosymbionts, often related to and likely derived from *Sodalis*, indicates their independent origin and convergence on supplementing insect hosts with deficient nutrients. These patterns seem to be repeated among diverse gammaproteobacteria infecting 57% of the surveyed planthoppers. Some of them seem to form relatively long-term associations with hosts, exemplified by *Purcelliella* in the family Cixiidae. Other associations may be more recent or even largely transient, as evidenced by symbiont genus-level distribution on the host phylogeny, and especially intra-species infection polymorphism revealed using 16S rRNA amplicon sequencing (Fig. 2). In many cases, however, it is impossible to conclude about the stability of the association based on the currently available data. McCutcheon et al. (2019) presented the challenges associated with phylogenetic reconstructions in these symbionts, whose genomes seem to undergo the spiral of rapid degeneration following the independent establishment in different hosts, with corresponding massive variations in evolutionary rates. Confident reconstruction of the evolutionary histories of these symbioses would require much more systematic sampling and genome-level datasets.

Associations of planthoppers with alphaproteobacterial symbionts seem to be less stable than symbiosis with Gammaproteobacteria. Observed in 35% of the surveyed planthopper species, these bacteria are not always fixed within populations. Facultative endosymbionts, the functional category that many strains of *Wolbachia* and *Rickettsia* are assigned to, are characterized by their patchy distribution across and within insect clades and species, variable prevalence within populations, and ability to occasionally transmit horizontally in addition to vertical transmission (Ahmed et al., 2016; Heath et al., 1999). However, in at least some cases, they have formed stable or even obligatory associations with hosts (Hosokawa et al., 2010). On the other hand, Acetobacteraceae live in a range of environments (Komagata et al., 2014), and the association of genus *Asaia* with flowers as well as guts of mosquitoes and other insects suggests its frequent environmental transmission (Bassene et al., 2020; Favia et al., 2007). This may not always be the case, given its highly specific endobacterial localization in *Trypetimorpha occidentalis*. This unique association of *Sulcia* with *Asaia*-like symbiont requires deeper analyses and will be the subject of a subsequent study.

Combined, the existing data make it clear that the impressive diversity of symbioses across planthoppers is a combination of their long-term co-diversification with hosts and frequent independent infections with a few clades of bacteria and fungi. Following such infections, likely often leading to the replacement of previously colonizing microbes, the stability of the association and it s subsequent evolution may vary. Our ongoing genomics work on microbes associated with these and other planthoppers could reveal the nature and biological significance of many of the symbiotic associations.

### 4.2. Conserved nature of established symbioses

In four Auchenorrhynchan superfamilies, beta- or alphaproteobacterial symbionts have established different tissue localizations relative to their co-symbiont *Sulcia*, also dividing nutritional responsibilities differently (Łukasik et al., 2018; Mao et al., 2018; McCutcheon and Moran, 2010; Michalik et al., 2021). In all Cicadomorpha, *Sulcia* and its co-symbiont generally occupy different regions of the same bacteriome; *Sulcia* colonizes the outer portion of the bacteriome, and its partner is localized in the bacteriome’s central part. In planthoppers, our microscopic survey across families separated by up to about 200 my of evolution revealed that *Sulcia* and *Vidania* always occupy separate bacteriomes. However, within a superfamily, as long as both ancestral symbionts are present, their nutritional functions and the organization of symbiont–containing tissue appear highly conserved (Buchner, 1965; Kobiałka et al., 2018; Koga et al., 2013; Łukasik et al., 2018; Michalik et al., 2018, 2021; Müller, 1940a, 1940b). As explained earlier, planthopper bacteriome size, shape, and localization within the abdominal cavity vary across species and between males and females of the same species. However, generally, there is no difference in the internal organization of bacteriome among sexes, as also reported from other auchenorrhynchans (Kobiałka et al., 2018; Szklarzewicz et al., 2020). The situation is different with the rectal organ, the second type of *Vidania* bacteriome found exclusively in females, which contain morphologically different *Vidania* cells. Given *Vidania* shape similarity to that observed during transovarial transmission, it has been proposed that this pool of *Vidania* cells is intended for the transmission to the progeny (Bressan and Mulligan, 2013; Buchner, 1965).

The only observed departures from the universal - ancestral organization were when other, more recently acquired symbionts established residence within *Sulcia* or *Vidania* bacteriomes. For example, in the family Cixiidae, *Vidania* always shares a common bacteriome with *Purcelliella*, whereas in *Pyrops clavatus, Sulcia* co-occurs with *Sodalis*. A unique example of such integration of a newly acquired symbiont in the biology of ancient one is *T. occidentalis*, where *Asaia*-like symbiont established residence within *Sulcia* cells. In the three planthopper clades characterized to date using genomics, different combinations of independently acquired, accessory symbionts seem to have had limited effects on the genomes or functions of the ancient symbionts. However, it remains to be demonstrated whether and how *Sulcia* and *Vidania* have been affected in some of the more complicated associations, including those described here.

### 4.3. Idiosyncracy in newly forming symbioses

Across our 44 planthopper species, we observed multiple types of the organization of tissues occupied by accessory symbionts representing different clades and derived from independent infections. Some of these distribution patterns closely resemble observations from other insects. For example, gammaproteobacterial symbiont *Arsenophonus* usually occupies distinct bacteriomes. We found this type of organization in 7 out of 8 planthopper species harboring this symbiont, and it was previously reported from other hemipterans, including leafhoppers, whiteflies, and scale insects (Gottlieb et al., 2008; Kobiałka et al., 2018; Michalik et al., 2018). Other organization types are very unusual, and perhaps unique to planthoppers. For example, in *Tettigometra sulphurea, Pectobacteria* share the common bacteriome with another accessory symbiont - *Sodalis*, creating a bacteriome with a structure never before reported from Auchenorrhyncha or, to our knowledge, any other insects.

Closely related microbes may settle within different tissues when colonizing different hosts. For example, in most planthoppers, bacteria *Sodalis* inhabit separate bacteriomes, but sometimes they colonize the same organs as other symbionts, including *Sulcia* or *Pectobacterium*. Likewise, *Ophiocordyceps* fungi may be localized in mycetomes, or alternatively, dispersed in fat body cells. Both these widely distributed microbial clades, when colonizing other insects, have established in a wide range of tissues. Indicatively, *Sodalis*, depending on the host, may be localized in the bacteriocytes, gut lumen, gut epithelium, milk gland, or even inside the cytoplasm of other symbiotic bacteria (Attardo et al., 2008; Husnik and McCutcheon, 2016; Masson et al., 2015; Michalik et al., 2021). Similarly, *Ophiocordyceps*, which replaced *Hodgkinia* in different clades of Japanese cicadas, established either within an epithelium of the shared *Sulcia-Hodgkinia* bacteriome (resulting in a shared *Ophiocordyceps-Sulcia* bacteriome), or in a new type of bacteriome while *Sulcia* remained on its own (Matsuura et al., 2018).

The symbiont localization may also be pre-determined to at least some extent by the host internal environment, which is likely to vary consistently among host clades. A great example of such pre-determination are mealybugs - where independently established gamma-symbionts across multiple host species always localized inside the cells of *Tremblaya* (Husnik and McCutcheon, 2016). In planthoppers, we need both broader sampling and robust phylogenomic tools before talking about differences among host clades in the tissues where independently acquired symbionts localize. Then, an important question is how this diversity of tissue localizations is shaped and what determines the localization and distribution of microbes newly acquired by different host clades. It is likely that the localization is a balance between pre-adaptations in different microbial clades or strains, pre-adaptations in different insect lineages, evolutionary processes in both symbionts and hosts, and likely, a good deal of chance. Following a new colonization, the microbe - likely a pathogen failing to display virulent phenotype, or a versatile opportunist, faces multiple challenges, the most important of which are avoiding insect’s immune system response and colonizing its tissues (Bright and Bulgheresi, 2010). The initial localization of a newly arrived symbiont may also be determined by the mode of infection, as it may need to pass through multiple tissues to colonize the host. However, following successful transmission to subsequent generations, it is likely to establish within a certain tissue. We think that in the longer term, that localization is unlikely to change spontaneously. However, over time, evolutionary processes acting both on the host and symbiont genomes may change the nature and organization of the tissue where the symbiont is localized.

Conversely, localization may determine symbiont biology and evolution, including long-term prospects of a newly established infection. For example, *Purcelliella*, a *Sodalis*-allied symbiont of the Cixiidae planthoppers, stands out from among related strains by its greatly reduced genome (Bennett and Mao, 2018) and presence in all surveyed members of the family - both suggestive of an unusually stable association. It is tempting to speculate that this stability is related to its unique localization in bacteriomes shared with *Vidania*.

### 4.4. The importance of a multi-pronged characterization of insect symbiosis diversity

The knowledge of the symbiont diversity in planthopper tissues is not new. However, we have gone a long way since the times of Müller (1940a, b) and Buchner (1965), reconstructing the co-diversification and replacement patterns in Auchenorrhynchan symbionts using relatively simple microscopy techniques, yet still with impressive accuracy. The rapid development of sequencing-based techniques has greatly simplified the task of characterizing host-symbiont associations. Through a combination of broad sampling and cost-effective screens, we can uncover the general patterns of symbiont diversity, distribution, and stability within clades. Shotgun metagenomics enables high-resolution phylogenomic reconstructions of host-symbiont relationships and informs us of putative symbiont functions. Transcriptomics and proteomics are powerful approaches to verify these functions. Finally, all these approaches can be combined with genetic manipulations and experiments to unequivocally demonstrate the nature and significance of broadly relevant processes (Bublitz et al., 2019; Su et al., 2022).

Unfortunately, such combinations of cutting-edge tools have not been applied to many systems, and outside of a few model organisms, knowledge about the diversity, distribution, evolutionary patterns, and biological significance of symbiotic microorganisms is lacking. Symbiont distribution within host tissues is also among these understudied areas, despite being critical for understanding the nature of host-symbiont interactions. The current study shows how comparative microscopy can complement increasingly popular sequencing-based approaches, in addressing a series of questions about how the symbioses are organized, how they function, and how they evolve. As we proceed in our attempts to describe the diversity and nature of host-microbial associations and their stability and roles in our rapidly changing world, microscopy and sequencing will form a particularly powerful combination of research tools.

## 5. Acknowledgements

We thank Ada Jankowska for laboratory assistance and help drafting figures, Monika Prus-Frankowska and Mateusz Buczek for laboratory assistance, and John McCutcheon for permission to use the facilities at the University of Montana. This project was supported by the Polish National Science Centre grants 2017/26/D/NZ8/00799 (to A.M.) and 2018/30/E/NZ8/00880 (to P.Ł.) and Polish National Agency for Academic Exchange grant PPN/PPO/2018/1/00015 (P.Ł.).

## Conflict of interests

The authors declare no conflict of interests

## Data availability

Sequence data have been deposited in GenBank under accession numbers provided in Supplementary Table S1 and at https://github.com/AnnaMichalik22/Bacteriome-organization-in-planthoppers.git

## Notes

### Competing Interest Statement

The authors have declared no competing interest.

### Summary of Updates

Figure 1 revised

## References

Ahmed, M.Z., Breinholt, J.W., Kawahara, A.Y., 2016. Evidence for common horizontal transmission of Wolbachia among butterflies and moths. BMC Evol. Biol. 16, 118. https://doi.org/10.1186/s12862-016-0660-x

Attardo, G.M., Lohs, C., Heddi, A., Alam, U.H., Yildirim, S., Aksoy, S., 2008. Analysis of milk gland structure and function in Glossina morsitans: Milk protein production, symbiont populations and fecundity. J. Insect Physiol. 54, 1236–1242. https://doi.org/10.1016/j.jinsphys.2008.06.008

Bassene, H., Niang, E.H.A., Fenollar, F., Doucoure, S., Faye, O., Raoult, D., Sokhna, C., Mediannikov, O., 2020. Role of plants in the transmission of Asaia sp., which potentially inhibit the Plasmodium sporogenic cycle in Anopheles mosquitoes. Sci. Rep. 10, 7144. https://doi.org/10.1038/s41598-020-64163-5

Baumann, P., 2005. Biology of bacteriocyte-associated endosymbionts of plant sap-sucking insects. Annu. Rev. Microbiol. 59, 155–189. https://doi.org/10.1146/annurev.micro.59.030804.121041

Bennett, G.M., Mao, M., 2018. Comparative genomics of a quadripartite symbiosis in a planthopper host reveals the origins and rearranged nutritional responsibilities of anciently diverged bacterial lineages. Environ. Microbiol. 20, 4461–4472. https://doi.org/10.1111/1462-2920.14367

Bennett, G.M., McCutcheon J. P., MacDonald B. R., Romanovicz D., Moran N. A., 2014. Differential genome evolution between companion symbionts in an insect-bacterial symbiosis. mBio 5, e01697–14. https://doi.org/10.1128/mBio.01697-14

Bennett, G.M., Moran, N.A., 2013. Small, smaller, smallest: the origins and evolution of ancient dual symbioses in a phloem-feeding insect. Genome Biol. Evol. 5, 1675–1688. https://doi.org/10.1093/gbe/evt118

Bourgoin, T., 2022. FLOW (Fulgoromorpha Lists On the Web): a world knowledge base dedicated to Fulgoromorpha.

Bressan, A., Mulligan, K.L., 2013. Localization and morphological variation of three bacteriome-inhabiting symbionts within a planthopper of the genus Oliarus (Hemiptera: Cixiidae). Environ. Microbiol. Rep. 5, 499–505. https://doi.org/10.1111/1758-2229.12051

Bright, M., Bulgheresi, S., 2010. A complex journey: transmission of microbial symbionts. Nat. Rev. Microbiol. 8, 218–230. https://doi.org/10.1038/nrmicro2262

Bublitz, D.C., Chadwick, G.L., Magyar, J.S., Sandoz, K.M., Brooks, D.M., Mesnage, S., Ladinsky, M.S., Garber, A.I., Bjorkman, P.J., Orphan, V.J., McCutcheon, J.P., 2019. Peptidoglycan production by an insect-bacterial mosaic. Cell 179, 703-712.e7. https://doi.org/10.1016/j.cell.2019.08.054

Buchner, P., 1965. Endosymbiosis of Animals with Plant Microorganisms. Interscience Publishers, New York.

Chan, P.P., Lin, B.Y., Mak, A.J., Lowe, T.M., 2021. tRNAscan-SE 2.0: improved detection and functional classification of transfer RNA genes. Nucleic Acids Res. 49, 9077–9096. https://doi.org/10.1093/nar/gkab688

Chernomor, O., von Haeseler, A., Minh, B.Q., 2016. Terrace aware data structure for phylogenomic inference from supermatrices. Syst. Biol. 65, 997–1008. https://doi.org/10.1093/sysbio/syw037

Deng, J., Bennett, G.M., Castillo Franco, D., Prus-Frankowska, M., Michalik, A., Łukasik, P., 2022. Genome comparison reveals inversions and alternative evolutionary history of nutritional endosymbionts in planthoppers (Hemiptera: Fulgoromorpha). In Prep.

Douglas, A.E., 2016. How multi-partner endosymbioses function. Nat. Rev. Microbiol. 14, 731–743.

Douglas, A.E., 2009. The microbial dimension in insect nutritional ecology. Funct. Ecol. 23, 38–47. https://doi.org/10.1111/j.1365-2435.2008.01442.x

Eddy, S.R., 2011. Accelerated profile HMM searches. PLOS Comput. Biol. 7, 1–16. https://doi.org/10.1371/journal.pcbi.1002195

Enomoto, S., Chari, A., Clayton, A.L., Dale, C., 2017. Quorum sensing attenuates virulence in Sodalis praecaptivus. Cell Host Microbe 21, 629-636.e5. https://doi.org/10.1016/j.chom.2017.04.003

Fan, H.-W., Noda, H., Xie, H.-Q., Suetsugu, Y., Zhu, Q.-H., Zhang, C.-X., 2015. Genomic analysis of an Ascomycete fungus from the rice planthopper reveals how it adapts to an endosymbiotic lifestyle. Genome Biol. Evol. 7, 2623–2634. https://doi.org/10.1093/gbe/evv169

Favia, G., Ricci, I., Damiani, C., Raddadi, N., Crotti, E., Marzorati, M., Rizzi, A., Urso, R., Brusetti, L., Borin, S., Mora, D., Scuppa, P., Pasqualini, L., Clementi, E., Genchi, M., Corona, S., Negri, I., Grandi, G., Alma, A., Kramer, L., Esposito, F., Bandi, C., Sacchi, L., Daffonchio, D., 2007. Bacteria of the genus Asaia stably associate with Anopheles stephensi, an Asian malarial mosquito vector. Proc. Natl. Acad. Sci. 104, 9047–9051. https://doi.org/10.1073/pnas.0610451104

Feldhaar, H., 2011. Bacterial symbionts as mediators of ecologically important traits of insect hosts. Ecol. Entomol. 36, 533–543. https://doi.org/10.1111/j.1365-2311.2011.01318.x

Flórez, L.V., Biedermann, P.H.W., Engl, T., Kaltenpoth, M., 2015. Defensive symbioses of animals with prokaryotic and eukaryotic microorganisms. Nat. Prod. Rep. 32, 904–936. https://doi.org/10.1039/C5NP00010F

Frago, E., Zytynska, S.E., Fatouros, N.E., 2020. Microbial symbionts of herbivorous species across the insect tree, in: Mechanisms Underlying Microbial Symbiosis. Academic Press Inc., pp. 111–159.

Gottlieb, Y., Ghanim, M., Gueguen, G., Kontsedalov, S., Vavre, F., Fleury, F., Zchori-Fein, E., 2008. Inherited intracellular ecosystem: symbiotic bacteria share bacteriocytes in whiteflies. FASEB J. 22, 2591–2599. https://doi.org/10.1096/fj.07-101162

Gruber-Vodicka, H.R., Seah, B.K.B., Pruesse, E., 2020. phyloFlash: Rapid small-subunit rRNA profiling and targeted assembly from metagenomes. mSystems 5, e00920–20. https://doi.org/10.1128/mSystems.00920-20

Hammer, T.J., Moran, N., 2019. Links between metamorphosis and symbiosis in holometabolous insects. Philos Trans R Soc Lond B Biol Sci 20190068.

Harris, H.L., Brennan, L.J., Keddie, B.A., Braig, H.R., 2010. Bacterial symbionts in insects: balancing life and death. Symbiosis 51, 37–53. https://doi.org/10.1007/s13199-010-0065-3

Heath, B.D., Butcher, R.D.J., Whitfield, W.G.F., Hubbard, S.F., 1999. Horizontal transfer of Wolbachia between phylogenetically distant insect species by a naturally occurring mechanism. Curr. Biol. 9, 313–316. https://doi.org/10.1016/S0960-9822(99)80139-0

Hosokawa, T., Koga, R., Kikuchi, Y., Meng, X.-Y., Fukatsu, T., 2010. Wolbachia as a bacteriocyte-associated nutritional mutualist. Proc. Natl. Acad. Sci. 107, 769. https://doi.org/10.1073/pnas.0911476107

Howard, F.W., Weissling, T.J., O’brien, L.B., 2001. The larval habitat of Cedusa inflata (Hemiptera: Auchenorrhyncha: Derbidae) and its relationship with adult distribution on palms. Fla. Entomol. 84, 119–122.

Husnik, F., McCutcheon, J.P., 2016. Repeated replacement of an intrabacterial symbiont in the tripartite nested mealybug symbiosis. Proc. Natl. Acad. Sci. 113, E5416. https://doi.org/10.1073/pnas.1603910113

Johnson, K.P., Dietrich, C.H., Friedrich, F., Beutel, R.G., Wipfler, B., Peters, R.S., Allen, J.M., Petersen, M., Donath, A., Walden, K.K.O., Kozlov, A.M., Podsiadlowski, L., Mayer, C., Meusemann, K., Vasilikopoulos, A., Waterhouse, R.M., Cameron, S.L., Weirauch, C., Swanson, D.R., Percy, D.M., Hardy, N.B., Terry, I., Liu, S., Zhou, X., Misof, B., Robertson, H.M., Yoshizawa, K., 2018. Phylogenomics and the evolution of hemipteroid insects. Proc. Natl. Acad. Sci. 115, 12775. https://doi.org/10.1073/pnas.1815820115

Kalyaanamoorthy, S., Quang Minh, B., Wong, T.K., von Haeseler, A., Jermiin, L.S., 2017. ModelFinder: fast model selection for accurate phylogenetic estimates. Nat. Methods 587–589.

Kobiałka, M., Michalik, A., Szwedo, J., Szklarzewicz, T., 2018. Diversity of symbiotic microbiota in Deltocephalinae leafhoppers (Insecta, Hemiptera, Cicadellidae). Arthropod Struct. Dev. 47, 268–278. https://doi.org/10.1016/j.asd.2018.03.005

Kobiałka, M., Michalik, A., Walczak, M., Junkiert, Ł., Szklarzewicz, T., 2016. Sulcia symbiont of the leafhopper Macrosteles laevis (Ribaut, 1927) (Insecta, Hemiptera, Cicadellidae: Deltocephalinae) harbors Arsenophonus bacteria. Protoplasma 253, 903–912. https://doi.org/10.1007/s00709-015-0854-x

Koga, R., Bennett, G.M., Cryan, J.R., Moran, N.A., 2013. Evolutionary replacement of obligate symbionts in an ancient and diverse insect lineage. Environ. Microbiol. 15, 2073–2081. https://doi.org/10.1111/1462-2920.12121

Komagata, K., Iino, T., Yamada, Y., 2014. The Family Acetobacteraceae, in: Rosenberg, E., DeLong, E.F., Lory, S., Stackebrandt, E., Thompson, F. (Eds.), The Prokaryotes: Alphaproteobacteria and Betaproteobacteria. Springer Berlin Heidelberg, Berlin, Heidelberg, pp. 3–78. https://doi.org/10.1007/978-3-642-30197-1_396

Li, D., Luo, R., Liu, C.-M., Leung, C.-M., Ting, H.-F., Sadakane, K., Yamashita, H., Lam, T.-W., 2016. MEGAHIT v1.0: A fast and scalable metagenome assembler driven by advanced methodologies and community practices. Methods 102, 3–11. https://doi.org/10.1016/j.ymeth.2016.02.020

Łukasik, P., Chong, R.A., Nazario, K., Matsuura, Y., Bublitz, D.A.C., Campbell, M.A., Meyer, M.C., Van Leuven, J.T., Pessacq, P., Veloso, C., Simon, C., McCutcheon, J.P., 2019. One hundred mitochondrial genomes of cicadas. J. Hered. 110, 247–256. https://doi.org/10.1093/jhered/esy068

Łukasik, P., Nazario, K., Van Leuven, J.T., Campbell, M.A., Meyer, M., Michalik, A., Pessacq, P., Simon, C., Veloso, C., McCutcheon, J.P., 2018. Multiple origins of interdependent endosymbiotic complexes in a genus of cicadas. Proc. Natl. Acad. Sci. 115, E226. https://doi.org/10.1073/pnas.1712321115

Mao, M., Yang, X., Bennett, G.M., 2018. Evolution of host support for two ancient bacterial symbionts with differentially degraded genomes in a leafhopper host. Proc. Natl. Acad. Sci. 115, E11691. https://doi.org/10.1073/pnas.1811932115

Masson, F., Moné, Y., Vigneron, A., Vallier, A., Parisot, N., Vincent-Monégat, C., Balmand, S., Carpentier, M.-C., Zaidman-Rémy, A., Heddi, A., 2015. Weevil endosymbiont dynamics is associated with a clamping of immunity. BMC Genomics 16, 819. https://doi.org/10.1186/s12864-015-2048-5

Matsuura, Y., Moriyama, M., Łukasik, P., Vanderpool, D., Tanahashi, M., Meng, X.-Y., McCutcheon, J.P., Fukatsu, T., 2018. Recurrent symbiont recruitment from fungal parasites in cicadas. Proc. Natl. Acad. Sci. 115, E5970. https://doi.org/10.1073/pnas.1803245115

McCutcheon, Boyd, Dale, 2019. The life of an insect endosymbiont from the cradle to the grave. Curr. Biol. 29, R485–R495. https://doi.org/10.1016/j.cub.2019.03.032

McCutcheon, McDonald, Moran, 2009. Convergent evolution of metabolic roles in bacterial co-symbionts of insects. Proc. Natl. Acad. Sci. 106, 15394–15399.

McCutcheon, Moran, 2010. Functional convergence in reduced genomes of bacterial symbionts spanning 200 My of evolution. Genome Biol. Evol. 2, 708–718. https://doi.org/10.1093/gbe/evq055

McFadden, G.I., 2001. Primary and secondary endosymbiosis and the origin of plastids. J. Phycol. 37, 951–959. https://doi.org/10.1046/j.1529-8817.2001.01126.x

Michalik, A., Castillo Franco, D., Kobiałka, M., Szklarzewicz, T., Stroiński, A., Łukasik, P., 2021. Alternative transmission patterns in independently acquired nutritional co-symbionts of Dictyopharidae planthoppers. mBio 12, e01228–21. https://doi.org/10.1101/2021.04.07.438848

Michalik, A., Jankowska, W., Kot, M., Gołas, A., Szklarzewicz, T., 2014. Symbiosis in the green leafhopper, Cicadella viridis (Hemiptera, Cicadellidae). Association in statu nascendi? Arthropod Struct. Dev. 43, 579–587. https://doi.org/10.1016/j.asd.2014.07.005

Michalik, A., Jankowska, W., Szklarzewicz, T., 2009. Ultrastructure and transovarial transmission of endosymbiotic microorganisms in Conomelus anceps and Metcalfa pruinosa (Insecta, Hemiptera, Fulgoromorpha). Folia Biol. Kraków 57, 131–137.

Michalik, A., Szwedo, J., Stroiński, A., Swierczewski, D., Szklarzewicz, T., 2018. Symbiotic cornucopia of the monophagous planthopper Ommatidiotus dissimilis (Fallén, 1806) (Hemiptera: Fulgoromorpha: Caliscelidae). Protoplasma 255, 1317–1329. https://doi.org/10.1007/s00709-018-1234-0

Miller, M.A., Pfeiffer, W., Schwartz, T., 2010. Creating the CIPRES Science Gateway for inference of large phylogenetic trees. Gatew. Comput. Environ. Workshop GCE 1–8. https://doi.org/10.1109/GCE.2010.5676129

Minh, B.Q., Schmidt, H.A., Chernomor, O., Schrempf, D., Woodhams, M.D., von Haeseler, A., Lanfear, R., 2020. IQ-TREE 2: New models and efficient methods for phylogenetic inference in the genomic era. Mol. Biol. Evol. 37, 1530–1534. https://doi.org/10.1093/molbev/msaa015

Moran, N.A., McCutcheon, J.P., Nakabachi, A., 2008. Genomics and evolution of heritable bacterial symbionts. Annu. Rev. Genet. 42, 165–190. https://doi.org/10.1146/annurev.genet.41.110306.130119

Moran, N.A., Munson, M.A., Baumann, P., Ishikawa, H., 1993. A molecular clock in endosymbiotic bacteria is calibrated using the insect hosts. Proc. R. Soc. Lond. B Biol. Sci. 253, 167–171. https://doi.org/10.1098/rspb.1993.0098

Moran, N.A., Tran, P., Gerardo, N.M., 2005. Symbiosis and insect diversification: an ancient symbiont of sap-feeding insects from the bacterial phylum Bacteroidetes. Appl. Environ. Microbiol. 71, 8802. https://doi.org/10.1128/AEM.71.12.8802-8810.2005

Müller, H.J., 1949. Zur Systematik und Phylogenie der Zikadenendosymbiose. Biol Zb 343–368.

Müller, H.J., 1940a. Die Symbiose der Fulgoroiden (Homoptera Cicadina). Zoologica 98, 1–110.

Müller, H.J., 1940b. Die Symbiose der Fulgoroiden (Homoptera-Cicadina). Zoologica, Die Symbiose der Fulgoroiden 98, 111–220.

Nishino, T., Tanahashi, M., Lin, C.-P., Koga, R., Fukatsu, T., 2016. Fungal and bacterial endosymbionts of eared leafhoppers of the subfamily Ledrinae (Hemiptera: Cicadellidae). Appl. Entomol. Zool. 51, 465–477. https://doi.org/10.1007/s13355-016-0422-7

Noda, H., Nakashima, N., Koizumi, M., 1995. Phylogenetic position of yeast-like symbiotes of rice planthoppers based on partial 18S rDNA Sequences. Insect Biochem. Mol. Biol. 25, 639–646. https://doi.org/10.1016/0965-1748(94)00107-S

Oliver, K.M., Degnan, P.H., Burke, G.R., Moran, N.A., 2010. Facultative symbionts in aphids and the horizontal transfer of ecologically important traits. Annu. Rev. Entomol. 55, 247–266. https://doi.org/10.1146/annurev-ento-112408-085305

Poole, A.M., Gribaldo, S., 2014. Eukaryotic Origins: How and when was the mitochondrion acquired? Cold Spring Harb Perspect Biol 6, a015990.

Song, N., Liang, A.-P., 2013. A preliminary molecular phylogeny of planthoppers (Hemiptera: Fulgoroidea) based on nuclear and mitochondrial DNA sequences. PLOS ONE 8, e58400. https://doi.org/10.1371/journal.pone.0058400

Su, Y., Lin, H.-C., Teh, L.S., Chevance, F., James, I., Mayfield, C., Golic, K.G., Gagnon, J.A., Rog, O., Dale, C., 2022. Rational engineering of a synthetic insect-bacterial mutualism. Curr. Biol. 32, 3925-3938.e6. https://doi.org/10.1016/j.cub.2022.07.036

Sudakaran, S., Kost, C., Kaltenpoth, M., 2017. Symbiont acquisition and replacement as a source of ecological innovation. Trends Microbiol. 25, 375–390. https://doi.org/10.1016/j.tim.2017.02.014

Szklarzewicz, T., Michalik, K., Grzywacz, B., Kalandyk-Kolodziejczyk, M., Michalik, A., 2021. Fungal associates of soft scale insects (Coccomorpha: Coccidae). Cells 10(8), 1922.

Szklarzewicz, T., Swierczewski, D., Stroiński, A., Michalik, A., 2020. Conservatism and stability of the symbiotic system of the invasive alien treehopper Stictocephala bisonia (Hemiptera, Cicadomorpha, Membracidae). Ecol. Entomol. 45, 876–885. https://doi.org/10.1111/een.12861

Urban, J.M., Cryan, J.R., 2012. Two ancient bacterial endosymbionts have coevolved with the planthoppers (Insecta: Hemiptera: Fulgoroidea). BMC Evol. Biol. 12, 87. https://doi.org/10.1186/1471-2148-12-87

Wheeler, T.J., Eddy, S.R., 2013. nhmmer: DNA homology search with profile HMMs. Bioinformatics 29, 2487–2489. https://doi.org/10.1093/bioinformatics/btt403

Wickham, H., 2011. ggplot2. WIREs Comput. Stat. 3, 180–185. https://doi.org/10.1002/wics.147

Wilson, S.W., O’Brien, L.B., 1987. Survey of planthopper pests of economically important plants (Homoptera: Fulgoroidea), in: Proceedings of 2nd International Workshop on Leafhoppers and Planthoppers of Economic Importance: Brigham Young University, Provo, Utah, USA, 28th July-1st August 1986/Edited by MR Wilson, LR Nault. London: CAB International Institute of Entomology, 1987.

Zytynska, S.E., Tighiouart, K., Frago, E., 2021. Benefits and costs of hosting facultative symbionts in plant-sucking insects: A meta-analysis. Mol. Ecol. 30, 2483–2494. https://doi.org/10.1111/mec.15897

